# Role of CYP3A5 in modulating androgen receptor signaling and its relevance to African American men with prostate cancer

**DOI:** 10.1101/2020.01.24.918920

**Authors:** Priyatham Gorjala, Oscar B Goodman, Rick A Kittles, Ranjana Mitra

## Abstract

**Background:** Androgen receptor signaling is crucial for prostate cancer growth and is regulated by intratumoral CYP3A5. As African American (AA) men often carry the wild type CYP3A5 and express high level of CYP3A5 protein, we tested the effect of blocking the wild type CYP3A5 in prostate cancer cells from AA men on androgen receptor signaling. CYP3A5 processes several commonly prescribed drugs and many of these are CYP3A5 inducers (e.g. phenytoin and rifampicin) or inhibitors (e.g. ritonavir and amiodarone). In this study, we test the effect of these commonly prescribed CYP3A5 inducers/inhibitors on AR signaling in prostate cancer cells.

**Methods:** Cell fractionation and immunofluorescence studies were performed to study AR nuclear localization and activation process using CYP3A5 siRNA and CYP3A5 inducers and inhibitors. A qPCR based array was employed to examine expression of AR downstream regulated genes after blocking CYP3A5 expression using a pool of CYP3A5 siRNA. Cell growth was monitored using MTS based assays. Since AAs tend to carry wild type CYP3A5 and non-Hispanic White Americans (NHWA) carry mutated CYP3A5 two cell lines one of AA origin (MDAPCA2b) carrying wt CYP3A5 and the other of NHWA origin (LNCaP) carrying mutant CYP3A5 were used for above experiments.

**Results:** Similar to that observed in LNCaP (mutant CYP3A5) earlier, CYP3A5 siRNA treated MDACPA2b (AA, wild type CYP3A5) cells showed decreased AR nuclear translocation and PSA production. q-PCR based profiler assay identified several AR regulated genes which were downregulated with CYP3A5 siRNA pool treatment performed with cDNA from CYP3A5 siRNA pool and NT treated MDAPCA2b cells. These downregulated genes include SCL45A3, FKBP5, NCAPD3, MYC, MME, ELL2, PIK3R3, HPRT1 and SPDEF with p-value of ≤0.005. These genes are known to regulate AR nuclear translocation, cell cycle progression and cell growth. SCL45A3, FKBP5, MYC, and ELL2 also showed decreased protein levels after CYP3A5 siRNA treatment.

Commonly prescribed drugs which are either CYP3A5 inhibitors (amiodarone, ritonavir) or inducers (phenytoin, rifampicin) were tested for their ability to alter AR signaling in both LNCaP and MDAPCa2b cells. The results show that the CYP3A5 inducers promoted AR nuclear translocation and downstream signaling whereas CYP3A5 inhibitors abrogated them. The increased nuclear AR observed with phenytoin and rifampicin (CYP3A inducers) treatment is abrogated in CYP3A5 siRNA treated MDAPCa2b cells, confirming that the activation of AR activity is specific to changes in CYP3A5 activity. Both the inducers tested demonstrated increased cell growth of prostate cancer cells, whereas the inhibitors showed reduced cell growth. The difference in growth is more pronounced in MDAPCa2b cells which carries a wild type CYP3A5 as compared to LNCaP with the exception of ritonavir which also downregulates total AR levels.

**Conclusions:** Concomitantly prescribed CYP3A5 modulating drugs may alter downstream AR signaling, cell growth and ADT efficacy in men, more so in AAs expressing wild type CYP3A5. Further, characterization and utilization of this observation how CYP3A5 inducers and inhibitors can alter AR signaling may provide guidance to physicians co-prescribing CYP3A5 modulating drugs to treat comorbidities in elderly patients undergoing ADT, particularly AA.

## Introduction

Androgen depletion therapy (ADT) is the standard treatment to manage castration-resistant prostate cancer (CRPC) [1]. Androgen receptor (AR) signaling changes; by gaining mutations and overexpressing to support growth, as the cancer becomes castration resistant. In all phases of prostate cancer AR remains active and is still expressed in patients undergoing ADT [2–4]. Mutated AR often can bypass the need for androgen activation, and can act as transcriptional activator in absence of androgens, promoting tumor growth [5]. Several new approaches are clinically available to block AR signaling one of them being blocking non-gonadal androgen synthesis [6]. Another useful approach is to disrupt tubulin dependent AR translocation to nucleus. Nonetheless, eventually the AR bypasses these strategies, leading to CRPC. It is highly therapeutically relevant to identify any additional mechanism to block AR nuclear translocation [7–9].

Our previous work shows that CYP3A5 has a critical role in AR signaling as it promotes AR nuclear translocation and downstream signaling promoting growth [10]. CYP3A5 is a cytochrome P450 enzyme primarily expressed in liver and small intestine. In liver, its main function is to process xenobiotics. CYP3A5 along with CYP3A4 metabolizes 50% of the commonly prescribed drugs [11]; these drugs are inducers, inhibitors and substrates of CYP3A enzymes. Apart from liver and small intestine CYP3A5 is also expressed in prostate where its normal function is to convert testosterone to its lesser active derivative, 6β-hydroxytestosterone [12]. Prostate cancer patients are typically elderly as the average age at diagnosis is 66 [13] and often suffer from comorbidities. Medications prescribed for these comorbidities can be inducers, inhibitors or substrates of CYP3A5 and hence can modify intratumoral CYP3A5 activity and alter AR signaling and response to ADT. Prior reports strongly support our hypothesis that therapeutic management of cancers is compromised by drug-induced expression of members of the CYP3A subfamily [14]. CYP3A5 is the main isoform expressed in prostate where as CYP3A4 is the most common isoform expressed in liver. Although CYP3A4 and CYP3A5 share 80% similarity they are differentially regulated [14–16]. CYP3A5 expressed in prostate is also differently regulated compared to the one expressed in liver as it has a 5’ UTR with androgen response elements (ARE).

Healthy prostate tissues are shown to express high basal levels of CYP3A5, but CYP3A5 expression in prostate cancer tissues is less well-characterized [17–19]. Different expression patterns in tumor cells may be due to polymorphic expression of CYP3A5. CYP3A5 has several variations most common being the CYP3A5*3, that carries a A>G mutation at position 6986 in the intron 3 (CYP3A5*3, rs776746 A>G). The presence of CYP3A5*3 results in aberrant splicing producing truncated non-functional protein [20]. 90% of the Caucasians carry the *3 mutation (*3/*3) whereas African Americans mostly carry (72%) the wild type CYP3A5 (*1/*1 or *1/*3) expressing the full-length protein.

Previously we demonstrated that full length CYP3A5 facilitates nuclear translocation of androgen receptor in prostate cancer cells [10]. We have also demonstrated that CYP3A5 specific inhibitor, azamulin, and siRNA-based knock down of CYP3A5 expression reduced AR nuclear translocation. In the current study, we investigated the effect of commonly co-prescribed CYP3A5 inhibitors /inducers with androgen deprivation therapy, which can alter AR nuclear translocation and its downstream signaling affecting treatment response. This study can provide a prescription guideline to clinicians prescribing the CYP3A5 modulators with ADT. As African American mostly express the full length CYP3A5 that promotes androgen receptor signaling and promotes prostate cancer growth, this study is very relevant to the AA patients that often have aggressive disease and present health disparity.

## Materials and Methods

### Cell lines, Drugs and antibodies

LNCaP, MDAPCa2b, 22RV1 and E066AAhT cells were purchased from ATCC and maintained in RPMI (Invitrogen, Carlsbad, CA), F-12K medium (ATCC® 30-2004), RPMI and DMEM (Invitrogen, Carlsbad, CA) media respectively. Supplements were added as recommended by ATCC. C4-2 was a gift from Dr. David Nanus and maintained in RPMI media. RC77 T/E (tumor) and RC77 N/E (normal) cell lines are a gift from Dr. Johng S Rhim and are maintained in Keratinocyte SFM media supplemented with epidermal growth factor and bovine pituitary extract [21].

Antibodies against Androgen receptor (ab74272), were obtained from Abcam, (Cambridge, MA). Anti-Flag (F1804) was from Millipore-Sigma (St. Louis, MO) and anti-GAPDH (10R-G109A) was from Fitzgerald Industries (Acton, MA). Anti-α tubulin (2125S) and anti Lamin A/C (4C11) were obtained from Cell signaling technologies (Danvers, MA). The secondary antibodies (IR dye 680 and IR dye 800) were from LI-COR (Lincoln, NE).

CYP3A inducers phenytoin (PHR1139), rifampicin (R3501), and inhibitors ritonavir (SML0491), amiodarone hydrochloride (A8423), azamulin (SML0485) were purchased from sigma-aldrich (St. Louis, MO).

The siRNA transfection reagents, oligofectamine and OPTIMEM were from Invitrogen; the siRNAs were from Dharmacon (Thermo scientific). MTS cell titer reagent was from Promega (Madison,WI).

### Western blotting

Cells were washed in phosphate buffer (PBS) and lysed in RIPA buffer (50 mM TIRS, 150 mM NaCl, 1% NP40, 0.5% sodium deoxycholate, 0.1% SDS, 2mM EDTA, 2mM EGTA) supplemented with protease and phosphatase inhibitors. GAPDH was used as an internal control for total protein, tubulin for cytoplasmic fraction and Lamin A/C for nuclear fraction. Infrared fluorescent-labeled secondary antibodies were used for detection using Odyssey CLx.

### Cell fractionation

Nuclear and cytoplasmic cell fractionation was prepared using NE-PER Nuclear and cytoplasmic extraction kit from Thermo Scientific (Cat no.78833) and manufacturer’s instructions were followed. The cells were treated with drugs in charcoal-stripped phenol red free media 48 hours after plating. Drugs and DHT were added at specified concentration and duration as indicated. Cells were washed in PBS once before cells were suspended in CER buffer. Protease, phosphatase inhibitors and EDTA was added prior to cell lysis. The pellet remaining after cytoplasmic isolation was washed twice with PBS. The pellet was suspended in NER buffer for nuclear fraction extraction according to guidelines, samples were stored at −80°C until further processing. Tubulin and Lamin were used as internal controls for cytoplasmic and nuclear fractions respectively.

### siRNA inhibition

Cells were plated in complete media without antibiotics on poly D-lysine-coated plates (80,000 cells per 6 well). After 48 hrs. of growth the cells were transfected using RNAimax according to the manufacturer’s instructions. The smart pool non-target (NT) siRNA (Dharmacon catalog# D-001810-10) was used as a transfection control with the experimental target gene siRNAs. A pool of four siRNA (Dharmacon catalog# L-009684-01) against the CYP3A5 were used to block the expression. The final concentration of the siRNA (NT and targets) used was 30 nM.

### Confocal Microscopy

Cells were seeded into 35 mm Glass bottom dish (Cellvis catalog# D35C4-20-1.5-N). The cells were fixed in 4% paraformaldehyde for 20 minutes and permeabilized using permeabilizing buffer (0.2% Tween 20 in PBS) for 5 minutes. Cells were blocked using 10% goat serum diluted in permeabilizing buffer with 1% BSA for 15 minutes. Primary antibodies were diluted at 1:100 in staining buffer (1% BSA in PBS) and incubated for 60 minutes at room temperature. Cells were washed three times (10 minutes each) in PBS. Secondary antibodies, Cy^5^-conjugated Donkey Anti-rabbit (711-175-152) and Alexa Fluor 488 – conjugated Donkey Anti-Mouse (715-545-150) from Jackson immuno research, West Grove, PA were diluted at 1:50 in staning buffer and incubated for 30 minutes at room temperature. Cells were washed three times (10 minutes) in PBS and stained with 1 µg/mL DAPI (4’, 6-diamidino-2-phenylindole) in PBS for 5 minutes at room temperature. The cells were stored in PBS at 4°C until imaging is completed.

Cells were imaged using confocal laser scanning microscopy on a Nikon A1R using a galvano scanner and a 60× Apo-TIRF oil immersion objective. To excite DAPI, FITC, TRITC and CY^5^ 405 nm, 488 nm, 561 nm and 638 nm solid-state lasers were used respectively. FITC and TRITC emissions were collected using GaAsP detectors on the A1R+ microscope. NIS Elements software form Nikon was used for recording the data.

### RT2 profiler

RNA was isolated using RNeasy Mini kit (Cat no.74104) from Qiagen (Germantown, MD) manufacturer’s instructions were followed. cDNA synthesis was performed using RT^2^ first strand kit (Cat no.330404) Qiagen, according to manufacturer’s instructions. The cDNA was diluted and used as template to analyze for gene expression pattern using RT^2^ Profiler PCR Arrays (Cat no. 330231) specifically designed to probe panel of Human Androgen Receptor Signaling Targets (PAHS-142Z). The real-time PCR reaction data was collected using ABI 7500 fast real-time PCR system. A total of 96 genes were profiled, data analysis was done using Geneglobe portal on Qiagen website. Samples (triplicates) were grouped into control (Non-Target) and test (CYP3A5 siRNA) and normalized with Beta-2-microglobulin (B2M) and Ribosomal protein, large, P0 (RPLP). A set of genes were identified based on fold change cutoff value of 2.0 and p value of 0.005.

### Luciferase assay

Cignal Lenti AR Reporter (luc) from Qiagen (product n0. 336851, Cat no. CLS-8019L) was used to generate AR pathway sensing LNCaP and MDAPCa2b cell lines for the study of the AR signal transduction pathway. These lentivirus particles have androgen response elements (ARE) fused to luciferase, which detects any changes in AR downstream signaling. Cells were transfected according to manufacturer instructions. Negative Control (only TATA box in place of ARE) transfected cell lines were also generated to measure background luciferase activity. Cells were maintained under puromycin selection pressure to select for stable chromosomal integration of the lentiviral constructs. The selected cells were tested for AR signaling pathway activation in response to DHT treatment after drug treatment using Brightglo luciferase assay from Promega (Cat no.E264A). The cells were collected into Eppendorf tube and divided into two equal aliquots. One aliquot was used for luciferase assay and was lysed with Glolysis buffer (Cat no. E266A) and the second aliquot was lysed with RIPA buffer for protein quantification.

### Genotyping Assay

DNA was isolated from cell lines using QIAam DNA mini Kit (Cat no. 51304) from Qiagen according to manufacturer instructions. TaqMan™ Drug Metabolism Genotyping Assay (Cat no. 4362691) from Applied Biosystems with 7500 Fast System was used to determine CYP3A5 *1 and *3 allelic status.

## RESULTS

### Differential expression of CYP3A5 between African American and Caucasian origin AR positive prostate cancer cell lines

We have previously shown that CYP3A5 is expressed in androgen receptor positive prostate cancer cell lines (LNCaP, C4-2 and 22RV1) and promotes activation of AR and prostate cancer growth [10]. CYP3A5 expression is polymorphic and is race linked so we genotyped the available AR positive cell lines from both African American (AA origin, MDAPCa2b, RC77 T/E (Tumor), RC77 N/E (normal) and Non-Hispanic White Americans (NHWA-LNCaP, C4-2, 22RV1, E006aahT) origin to determine their CYP3A5 polymorphism. (Table 1). Genotyping revealed that all the NHWA lines carry the *3/*3 CYP3A5 variant in homozygous form. The three cell lines from AA origin, carry *1/*3 heterozygous wild type/mutant CYP3A5. E006aahT has been found to be not of African American origin [22] and carries *3/*3 homozygous mutation. We used LNCaP (*3/*3) and MDAPCa2b (*1/*3) cells for our current study as they are of NHWA and AA origin respectively, and are AR positive commercially available (ATCC) and show similar response to androgens.

**Table 1:** CYP3A5 polymorphism analysis of commonly used prostate cancer cell lines. Seven androgen responsive prostate cell lines were tested for presence of wild type (*1) or mutant inactive (*3) CYP3A5 polymorphism by using a qPCR based genotyping assay.

| Cell Line | Genotype | Origin |
| --- | --- | --- |
| LNCaP | *3/*3 | NHWA |
| 22RV1 | *3/*3 | NHWA |
| C4-2 | *3/*3 | NHWA |
| E006AAhT | *3/*3 | NHWA |
| MDAPCa2b | *1/*3 | AA |
| RC77 T/E Tumor | *1/*3 | AA |
| RC77 N/E Normal | *1/*3 | AA |

### CYP3A5siRNA downregulates AR nuclear translocation in MDAPCa2b cells expressing wild type CYP3A5 (*1/*3)

To test if wild type full length CYP3A5 regulates AR nuclear activation in a similar fashion we used MDAPCa2b cells, which express wild type CYP3A5 (*1/*3). MDAPCa2b cells were treated with NT and CYP3A5 siRNA pool to specifically block CYP3A5 and then induced with DHT in charcoal stripped phenol red free media to monitor AR nuclear translocation and activation. Cells were then stained with AR and Cy5 labelled secondary antibody. CYP3A5 siRNA treatment resulted in decreased nuclear translocation of AR (Fig. 1A). This observation was further confirmed with cell fractionation experiments performed after NT and CYP3A5 siRNA treatment and DHT induction. Western blotting analysis was performed to monitor AR nuclear translocation in the cytoplasmic and nuclear fractions. The result confirms our previous observation with LNCaP cell line, CYP3A5 siRNA treated MDAPCA2b cells show decreased nuclear translocation of AR (Fig. 1C) after DHT induction as compared to non-target siRNA pool treated cells. In the NT siRNA treated group we observed significant cytoplasmic surge which was absent in the CYP3A5 siRNA treated cells. Of note CYP3A5 siRNA treatment did not affect total AR protein expression (Fig 1B).

**Figure 1:**
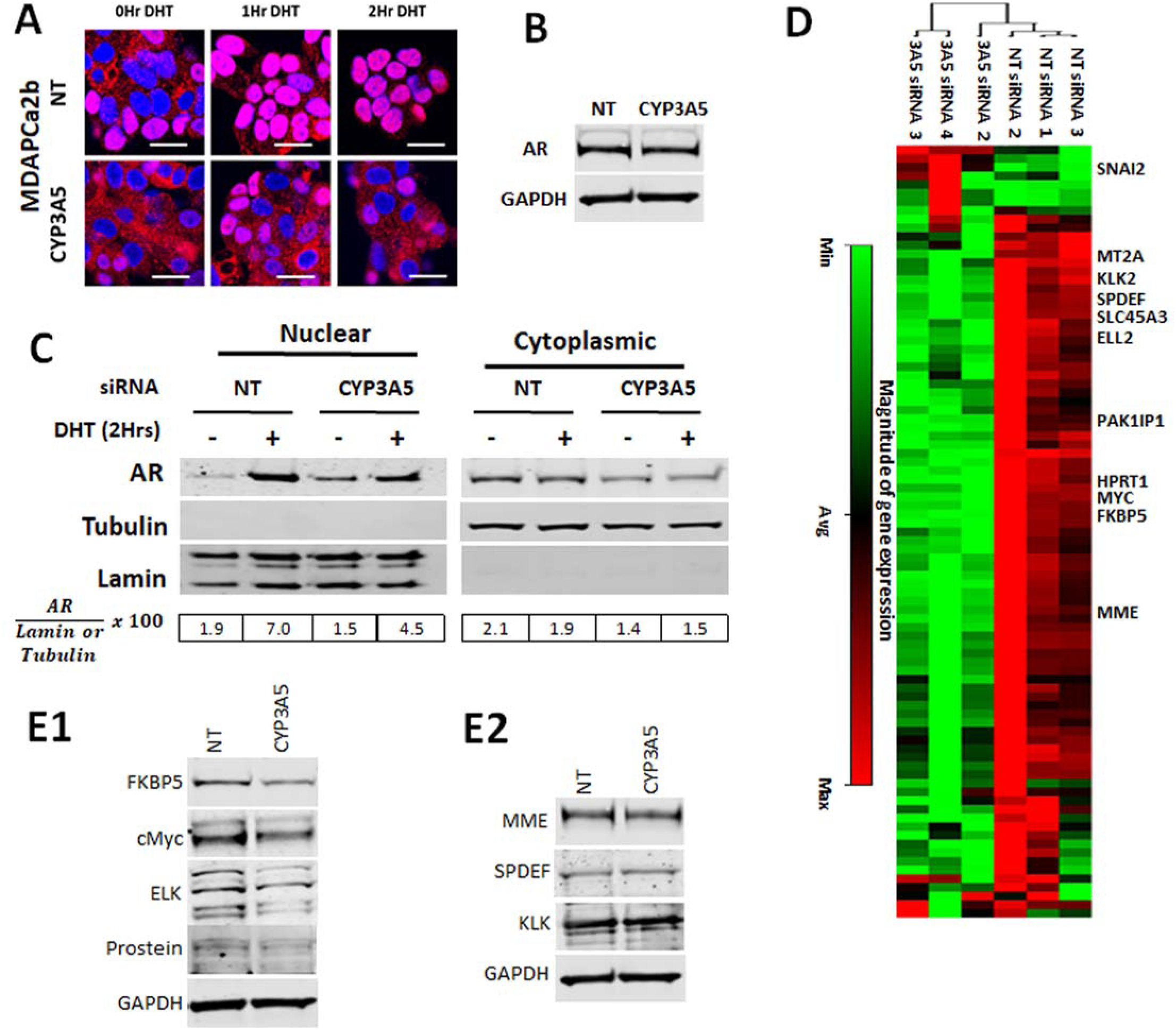
CYP3A5 siRNA downregulates AR nuclear translocation and expression of AR downstream regulated genes: **(A, C)** MDAPCa2b cells were transfected with CYP3A5 and non target (NT) siRNA. After 72 hours the cells were given 10nM DHT treatment (0, 1 and 2 hours). For microscopy **(A)** the cells were labelled with AR primary antibody and Cy5 secondary. The scale bar represents 50 µm. **(B)** Total protein was used to monitor changes in total AR protein expression. **(C)** After cell fractionation, the western blotting was performed using cytoplasmic and nuclear fractions and probed for AR, Tubulin and Lamin. **(D)** MDAPCa2b cells were seeded into 6 well plates. After 48 hours of allowing them to settle, the cells were treated with CYP3A5 siRNA or Non-Target siRNA and incubated for another 72 hours. RNA was isolated followed by cDNA preparation which was used in RT^2^ profiler assay. Fold-change values greater than 1 are indicated as positive- or an up-regulation (red) and less than −1 are indicated as negative or down-regulation (green). The P values are calculated based on a Student’s t-test of the replicate 2^ (- Delta CT) values for each gene in the control group and treatment groups. **(E1, E2)** Protein expression of the 7 genes that changes in gene expression was evaluated using western blotting. FKBP5, cMYC, ELK, Prostein (SLC45A3) showed decreased protein expression in response to CYP3A5 knock down. Whereas, MME, SPDEF and KLK showed no change.

### CYP3A5 siRNA downregulates expression of AR regulated genes in MDAPCA2b cells

To further evaluate the downstream signaling effect of CYP3A5 knockdown, cDNA prepared with RNA extracted from MDAPCA2b cells treated with NT and CYP3A5 siRNA pool was used for gene expression analysis. RT^2^ PCR pathway array deciphering changes in signaling targets downstream of Androgen receptor shows down regulation of several genes listed in Table 2 with fold changes greater than 2.0 and p-values less than 0.005 depicted (Fig. 1D). Western analysis was performed to evaluate whether gene expression changes translated into changes in protein expression. FKBP5, c-Myc, ELK-1, prostein protein expression also decreased in response to CYP3A5 siRNA treatment (Fig. 1E1) consistent with mRNA downregulation (fold change 0.68, 0.55, 0.49 and 0.45 respectively, P value ≤ 0.05). We did not observe changes in the levels of MME, SPDEF and KLK2 protein levels with CYP3A5 siRNA treatment (Fig. 1E2).

**Table 2:** CYP3A5 inhibition downregulates AR downstream regulated genes. Table showing down regulation of AR downstream genes with CYP3A5 siRNA treatment. These indicate that CYP3A5 activity modulating medications can affect AR downstream signaling regulating growth.

| AR Pathway Array |  |  |
| --- | --- | --- |
| Gene Symbol | Fold regulation | P value |
| SLC45A3 | -4.56 | 0.002 |
| FKBP5 | -4.43 | 0.002 |
| MYC | -3.68 | 0.001 |
| MME | -3.34 | 0.016 |
| PAK1IP1 | -3.25 | 0.016 |
| ELL2 | -3.25 | 0.004 |
| KLK2 | -2.82 | 0.009 |
| HPRT1 | -2.65 | 0.005 |
| SPDEF | -2.58 | 0.012 |
| MT2A | -2.45 | 0.001 |
| SNAI2 | 3.32 | 0.005 |

### Commonly co-prescribed CYP3A5 inducers /inhibitors can alter AR nuclear translocation

The average age at detection is 66 for prostate cancer patients; hence, they often have comorbidities, and are prescribed other medications to treat these co-morbidities while undergoing androgen deprivation treatment (ADT). CYP3A5 is known to process 33% of the commonly prescribed drugs and these co-prescribed drugs can be inducer/inhibitor of CYP3A5. Since AR is central to prostate cancer progression and is a main therapeutic target in treating prostate cancer any alteration in AR signaling can alter efficacy of these regimens. Based on our observation that CYP3A5 alters AR activity we wanted to test the effect of CYP3A5 inducers/inhibitors drugs on AR signaling as CYP3A5 can be modulated by these co-prescribed drugs. To evaluate the effect of known CYP3A inducers and inhibitors on AR nuclear translocation and downstream signaling we used two CYP3A inhibitors, amiodarone (5 µM) and ritonavir (35 µM); and two inducers, phenytoin (50 µM) and rifampicin (30 µg/mL) [23]. Amiodarone is often prescribed as an anti-arrhythmic drug whereas ritonavir is a component of highly active anti-retroviral therapy used in treating HIV patients. Phenytoin is one of the widely prescribed antiepileptic drugs and rifampicin is a known CYP3A5 inducer. CYP3A5 is not the main target of any of these drugs but its activity can be significantly modulated by these drugs. We tested their ability to affect AR activation process due to their ability to modulate CYP3A5 expression, which is separate from their primary target. Azamulin, a specific CYP3A inhibitor has been used as a control.

We tested the effect of these CYP3A5 inhibitors and inducers on total AR expression. Total cell lysates prepared from LNCaP and MDAPca2b cells incubated with the selected drugs were analyzed by western blotting to verify if protein expression of AR is affected. None of the drugs tested affects total AR protein expression in LNCaP and MDAPCa2b cells except ritonavir (Fig. 2A).

**Figure 2:**
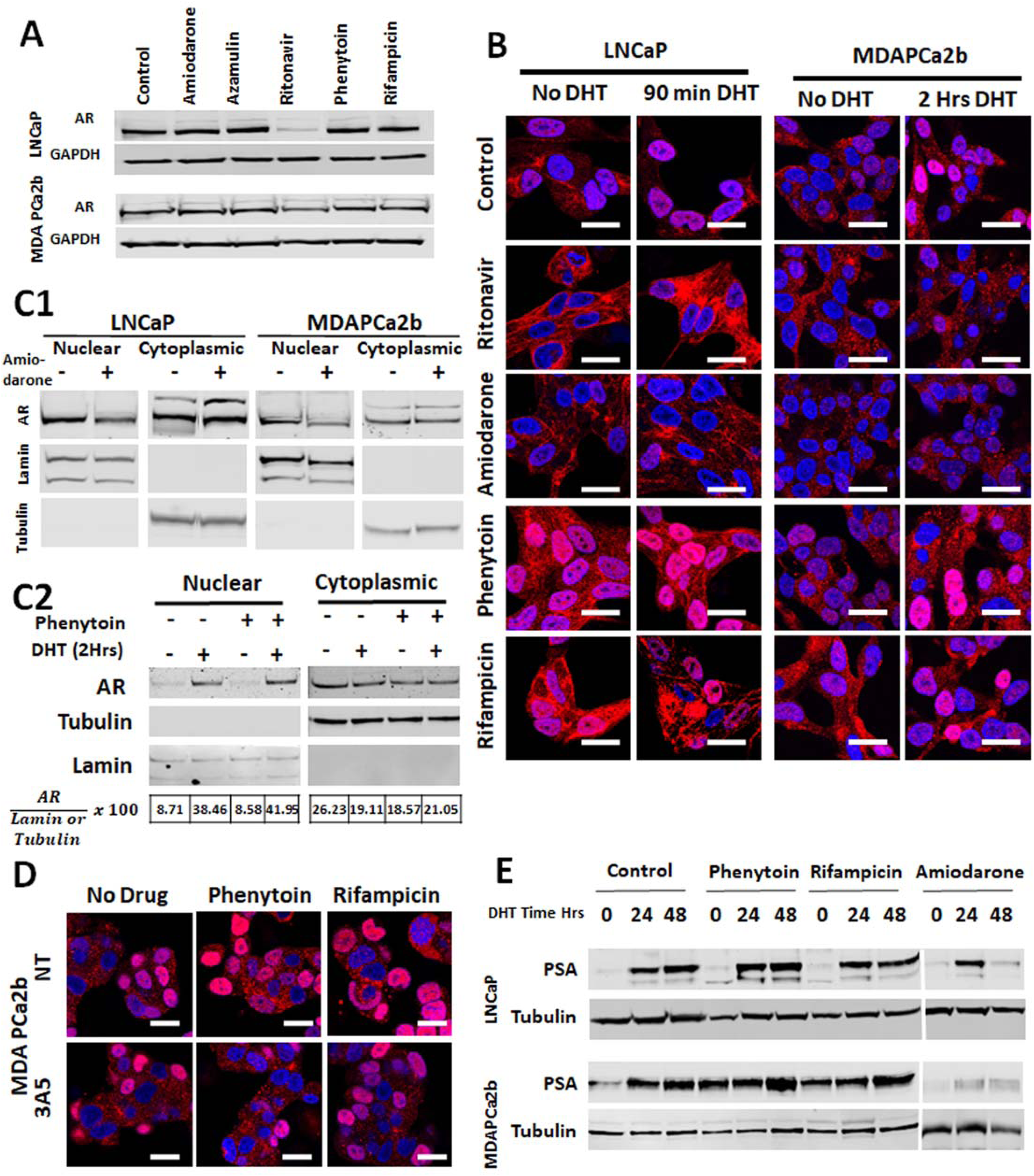
CYP3A5 inhibitors and inducers affect nuclear translocation of AR and its downstream signaling. **(A)** Effect of CYP inhibiting or inducing drugs on total AR expression. Total cell lysates from LNCaP and MDAPCa2b cells treated with CYP inhibitors (Ritonavir-35 µM, Azamulin 10 µM and Amiodarone-5 µM) and inducers (Phenytoin-50 µM and Rifampicin-30µg/ml) for 48 hours was used for western analysis. **(B)** Nuclear localization of AR after CYP3A5 inhibitor / inducer treatment. Immunostaining was performed on LNCaP and MDAPCa2b cells that were treated with CYP3A inhibitors (Ritonavir-35 µM and Amiodarone-5 µM) and inducers (Phenytoin-50 µM and Rifampicin-30µg/ml) for 48 hours in charcoal stripped serum media followed by with and without DHT induction (90min for LNCaP or 120 min for MDAPCa2b). Nucleus is stained with DAPI (blue), AR is stained with Cy5-secondary (red) antibody. Scale bar represents 25 µm. A section from center of z-stack is shown here to demonstrate the localization of AR in nucleus after treatments. **(C1)** Cell fractionation was performed after treating LNCaP and MDAPCa2b cells with Amiodarone (5µM) for 72 hours. The cytoplasmic and nuclear fractions were evaluated using western. **(C2)** MDAPCa2b cells were treated with phenytoin (50 µM) followed by 10nM DHT induction (120 min), nuclear and cytoplasmic fractions were analyzed by western blotting. Lamin and tubulin are controls for nuclear and cytoplasmic fractions. **(D)** AR nuclear translocation by CYP inducers in NT/CYP3A5 siRNA treated MDAPCa2b cells. MDAPCa2b cells were treated with NT/CYP3A5 siRNA for 24 hours and then incubated with CYP3A inducers, phenytoin (75µM) and rifampicin (30 µg/mL) for 48 hours. Confocal microscopy was performed and center of Z-stack is shown for nuclear AR localization. AR-red (Cy5) and nucleus (blue). Scale bar represents 25 µm. **(E)** To Confirm the effect of CYP3A5 modulating drugs on AR downstream signaling LNCaP and MDAPCa2b cells were treated with Phenytoin (50 µM), Rifampicin (30 µg/mL), Amiodarone (5 µM) in charcoal stripped serum media followed by 24 or 48 hours of DHT treatment. Total cell lysate was used to check PSA production using western analysis.

We observed reduced AR nuclear translocation in the cells treated with CYP3A inhibitors (amiodarone and ritonavir) and increased AR translocation in cells treated with CYP3A inducers (phenytoin and rifampicin) as compared to control cells that received no drugs (vehicle treated) (Fig. 2B) in both LNCaP and MDAPCa2b cell lines expressing different levels of CYP3A5 full length protein. Additionally, the cells treated with CYP3A inducers showed increase nuclear AR even without DHT induction compared to control. Similarly, the CYP3A5 inhibitor treated cells show lower nuclear AR also without DHT induction in both LNCaP (*3/*3) and MDAPCA2b (*1/*3) cells. We confirmed the CYP3A5 modulating effect of these drugs on AR activation by performing cell fractionation studies with and without DHT induction.

Nuclear AR is decreased in LNCaP and MDAPCa2b cells treated with amiodarone compared to control cells that did not receive drug treatment (Fig. 2C1). Similarly, phenytoin increased nuclear AR after DHT induction (Fig 2C2) in MDAPCa2b cells consistent with Immuno-florescence data. These observations indicate that changes in CYP3A5 levels in both LNCaP and MDAPCa2b cell lines can alter AR activation. Notably the small absolute changes in CYP3A5 levels observed in LNCaP cells expressing very low level of full length CYP3A5 can alter AR signaling significantly.

### Changes in AR activation by CYP3A inducers are due to their effect on CYP3A5 activity

To test our hypothesis that modulation in AR activation is dependent on the changes in CYP3A5 expression caused by the CYP3A5 inducers/inhibitors we performed AR nuclear localization assays after CYP3A5 and NT siRNA treatment. The MDAPCa2b cells treated with CYP3A5 siRNA and CYP3A5 inducer (phenytoin and rifampicin) do not show increased nuclear AR in contrast to NT control (Fig. 2D). This result supports that the observed changes in AR nuclear fraction is dependent on the modulation of CYP3A5 by the above mentioned CYP3A5 inducers (rifampicin and phenytoin) and is independent of their effect on main primary target.

### CYP3A5 inhibitors and inducers alter PSA levels

Prostate specific antigen (PSA) expression is regulated by androgen receptor and is an established marker to monitor AR downstream signaling. To evaluate downstream effects of AR nuclear translocation caused due to modulation of CYP3A5 we analyzed the level of PSA protein expression in the phenytoin, rifampicin and amiodarone treated cells. In both LNCaP and MDAPCa2b cell lines CYP3A inducing drugs phenytoin and rifampicin increased expression of PSA. The fold change in the MDAPCa2b cell line which carries a wild type CYP3A5 (Fig. 2E) was more as compared to LNCaP line carrying mutant CYP3A5 (*3/*3). As expected amiodarone reduced PSA protein expression in both the cell lines, the effect is more prominent after 48 hours of DHT treatment.

### CYP3A5 modulating drugs affect AR downstream signalling

We used a luciferase-based reporter assay to determine the effect of commonly prescribed CYP3A5 inhibitors/ inducers on their ability to modify AR downstream signaling. Both LNCaP and MDAPCa2b cell lines were transduced with a viral construct carrying androgen response elements (AREs) fused with luciferase, positive clones were selected after antibiotic selection. Negative controls were setup with constructs carrying only the TATA promoter without ARE. A pool of positive clones were used to monitor changes in luciferase activity after treatment with the CYP3A5 inducers/inhibitors. The reporter assay using MDAPCA2b cells show increased luciferase activity with CYP3A5 inducers (phenytoin, rifampicin and hyperforin) and decreased luciferase activity with inhibitors (ritonavir, amiodarone and chloramphenicol) (less AR activation) (Fig 3A). LNCaP cells showed increased luciferase activity after treatment with CYP3A5 inducers phenytoin and rifampicin with DHT treatment, phenytoin shows an increase in AR activity even without DHT induction similar to earlier observation (Fig 3B). The inhibitors amiodarone and ritonavir show reduced luciferase units (AR activity) both with or without DHT induction (Fig 3B).

**Figure 3:**
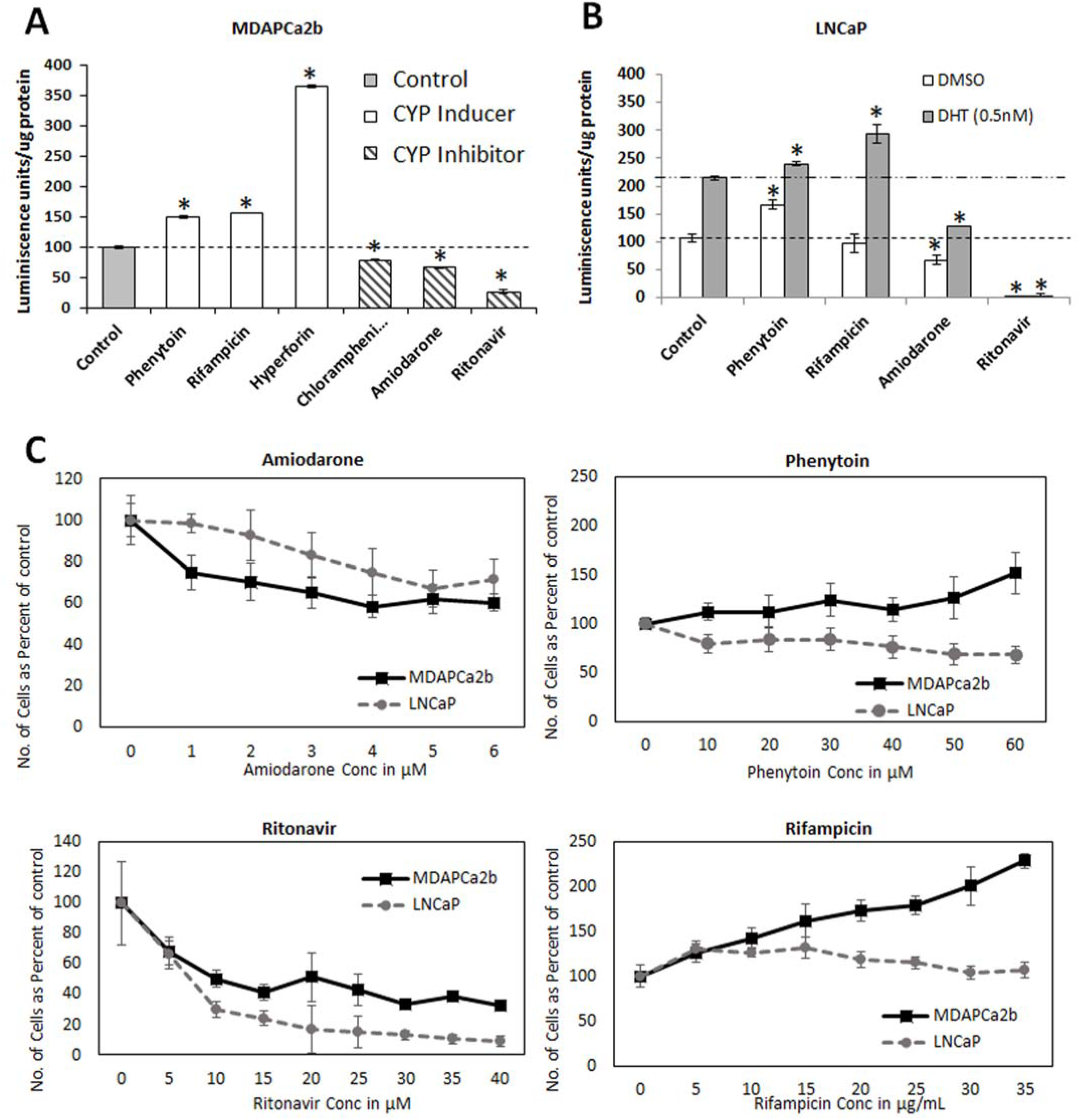
Effect of CYP inducers and inhibitors on cell growth in LNCaP and MDAPCa2b cells. **(A-B)** MDAPCa2b and LNCaP cells transfected with androgen response elements (ARE) fused to luciferase were used to evaluate AR downstream signaling activity. MDAPCa2b cells were treated with known CYP3A5 inducers (Phenytoin-50 µM, hyperforin-200μg/ml and Rifampicin-30µg/ml) and inhibitors (Ritonavir-35 µM, Azamulin-10 µM, chloramphenicol-10µM and Amiodarone-5 µM). LNCaP cells were treated with CYP inducing and inhibiting drugs in charcoal stripped serum followed by DHT (10nM) induction for one hour. In both cases CYP inducers showed increased AR signaling activity where as CYP inhibitors showed decreased AR signaling activity. **(C)** LNCaP and MDAPCa2b cells were treated with a CYP inhibitors, Amiodarone (0-6 µM) and Ritonavir (0-40 µM), CYP inducers Phenytoin (0-60 µM) and Rifampicin (0-35 µM) for 96 hours. The cell growth was accessed using MTS assay.

### CYP3A can regulate PCa cell growth by modifying AR activation

Androgen signaling pathway is involved in cell growth; based on our observation that CYP3A inhibitors and inducers alter AR nuclear translocation, we hypothesized that they should also alter cancer cell growth. To test our hypothesis, we monitored the effect of these inhibitors and inducers on prostate cancer cell growth. Both LNCaP and MDAPCa2b cell lines were incubated with different dose range of inducers [phenytoin (0-60 µM), rifampicin (0-35 µM)] and CYP3A inhibitors [amiodarone (0-6 µM), ritonavir (0-40 µM)]. Our results indicate that CYP3A inhibitors amiodarone and ritonavir decreased cell growth where as CYP3A inducers phenytoin and rifampicin reduce cell growth of both cell lines increasing concentrations (Fig 3C). The effect of CYP3A inducers and inhibitors are more pronounced in MDAPCa2b cells compared to LNCaP which may be due to the presence of wild type CYP3A5 (*1/*3) which has 3-4 times more functional CYP3A5 as compared LNCaP (*3/*3).

## DISCUSSION

Our previous work shows that CYP3A5 inhibition can lead to growth inhibition in LNCaP cells due to blocking of AR activation and downstream signaling. In keeping with previously published results for LNCaP, the MDAPCa2b, which carries one copy of wild type CYP3A5 (*1), also promotes AR nuclear localization. MDAPCa2b cells showed reduced AR nuclear localization after CYP3A5 siRNA (pool) treatment as compared to NT (pool). CYP3A5 is polymorphic with the wild type variant encoding full length translated protein being expressed in 73% of AAs, whereas only 5% of this variant is expressed in NHWA [20, 24]. Since *3 is the most common difference between AA and NHWA we analyzed the available prostate cancer cell lines and used one (*3/*3, LNCaP) and the other (*1/*3, MDAPCa2b) cell line for this study. There are 12 known SNPs in the CYP3A5 gene that mostly result in inactive protein. Distribution of these SNPs between races varies depending on the SNPs. The most commonly expressed mutation (*3) is a point mutation at 6986A > G that results in alternative splicing of an insertion from intron 3 resulting in a nonsense-mutated nonfunctional truncated protein. Even though A>G mutation leads to truncated protein in *3 mutation, 5% of the matured RNA can bypass the alternative splicing and express low levels of full length CYP3A5 protein as observed in LNCaP cells (*3/*3). The most common SNP (*3) mutation is present in 95% of NHWA, whereas 75% of AA carry wild type and 10-13% of AAs carry *6 and *7 mutations (truncated protein) [25, 26]. Prevalent expression of wild type CYP3A5 (*1/*1) form can promote AR activation in the AA prostate cancer patients as compared to NHWA. Since CYP3A5 is the major extrahepatic CYP3A isoform expressed in prostate and regulates AR activation, the presence of these SNPs in CYP3A5 may alter prostate cancer occurrence growth and treatment resistance in a race dependent manner.

Since MDAPCa2b carries a wt CYP3A5 we used this cell line for the PCR based pathway array to study the effect of CYP3A5 inhibition on AR downstream signaling. All the genes in this q-PCR base array carry androgen response elements and hence regulated by AR. The 11 genes which show maximum fold change (≥ 2.5) with CYP3A5 siRNA treatment are known to play important role in prostate cancer growth and severity. Further investigation revealed that four (SLC45A3, FKBP5, c-MYC and ELL2) of those 11 genes show reduced protein expression whereas MME, SPDEF and KLK2 did not show fold changes in protein expression. SLC45A3 also known as prostein is down regulated (−4.56 fold) with CYP3A5 siRNA treatment and belongs to solute carrier family 45. Protein expression is seen in both normal and malignant prostate tissue, its messenger RNA and protein are upregulated in response to androgen treatment in prostate cancer cells. [27, 28]. FKBP5 (downregulated, −4.43 fold, also called FKBP51) is a co-chaperone that belongs to a family of immunophilins, FK506 binding proteins (FKBPs). FKBP5 works with several different signaling pathways, including steroid receptor signaling, NF-kB, and AKT pathways, all of which contribute to tumorigenesis and drug resistance [29, 30] and FKBP5 is a target for AR signaling [31]. A recent study uncovered a mechanism in which FKBP5 is found to form a complex with HSP90 and promote AR signaling in prostate cancer [32]. Members of this family are targets for drugs such as rapamycin and cyclosporine. FKBP5 is known to modulate steroid receptor (androgen, progesterone, glucocorticoid) function by forming complex with HSP90 and HSP70. c-MYC also significantly downregulated with CYP3A5 siRNA treatment is one of the key genes amplified in prostate cancer progression. c-MYC induces AR gene transcription and is frequently upregulated in CRPC. A positive correlation between c-MYC and AR mRNA has been reported [33–37]. ELL2 (elongation factor, RNA polymerase II) is encoded by an androgen-response gene in the prostate [31, 38], it suppresses transient pausing of RNA polymerase II activity along the DNA strand and facilitates the transcription process [39]. ELL2 has been identified as an androgen response gene in immortalized normal human prostate epithelial cells as well as prostate cancer cell lines LNCaP and C4-2[31, 40]. ELL2 down regulation is seen in prostate cancer specimens and other observations indicate that its decrease improves cell proliferation, migration and invasion [41]. However, another study by Zang et. al. indicates that ELL2 has important role in DNA damage response and repair. This enables ELL2 loss to function like a double edge sword where on one hand can induce prostate carcinogenesis and on the other can sensitize cells to radiation therapy [42]. Human kallikrein-related peptidase 2 (KLK2, previously known as hK2) is a secreted serine protease from the same gene family as PSA. It shares 80% sequence homology with PSA and is responsible for cleavage of pre-PSA to active mature PSA [43]. Our studies only indicate fold change in mRNA levels but not protein levels after CYP3A5 siRNA treatment. Studies give contradictory evidence towards KLK2’s use as marker for detection of prostate cancer in combination with PSA [44–46]. KLK2 has been found to modulate AR to increase cell growth after development of CRPC [47]. In conclusion, this data supports our earlier observation that CYP3A5 plays a major role in AR regulation thus modulating AR downstream signaling and prostate cancer growth. This also points how presence of a wild type CYP3A5 (preferentially present in AAs) can significantly alter AR signaling compared to cells carrying only inactive CYP3A5 polymorphic forms (expressed in NHWAs).

CYP3A5 is an enzyme whose activity can be physiologically altered by many drugs that are activators or inhibitors of CYP3A5. Men with prostate cancer undergoing ADT are often elderly and have comorbidities requiring concomitant prescription medications, many of which are CYP3A5 inducers or inhibitors. The modulation of CYP3A5 by concomitant drugs may enhance or interfere with ADT, of great relevance to the AAs expressing wild type CYP3A5. Our data show that commonly prescribed CYP3A5 inducers promote AR nuclear migration whereas CYP3A5 inhibitors block AR nuclear migration. In our current study, we have used two CYP3A5 inhibitors (ritonavir and amiodarone) and two inducers (phenytoin and rifampicin) to test their effect on AR activation and downstream signaling in both LNCaP and MDAPCa2b cell lines. The results of the study indicate that the CYP3A5 inhibitors show less nuclear AR and less PSA expression similar to CYP3A5 siRNA. Whereas, the inducers on the other hand promoted nuclear AR translocation with and without DHT induction. Both the cell lines show similar effect since both the cell lines have different AR and CYP3A5 expression we were not able to derive a quantitative difference between both the cell lines. The reporter assay also showed similar response, which contains androgen response elements (AREs) to test the effect on AR downstream signaling. Although CYP3A5 is not the main target of any of these drugs it was shown that the specific effect on AR signaling is due to the changes in CYP3A5 and not due to the primary target of these drugs (Figure 2D).These drugs inhibit both CYP3A4 and CYP3A5 isoform. Since CYP3A5 is the major extrahepatic form expressed in prostate, the observed effect on AR signaling is due to the alteration in CYP3A5 and not CYP3A4.

Both the inducers increase the proliferation of the cells (LNCaP and MDAPCa2b) and the inhibitors reduce cell growth. Interestingly the effect of inducers and inhibitors on growth are more pronounced in MDAPCa2b which carries the wild type CYP3A5 as compared to LNCaP(*3/*3), with the exception of ritonavir. The observed difference can be because the other three tested inhibitors only effect the AR nuclear localization whereas ritonavir also affects total AR levels. Nonetheless, these data strongly suggest that concomitant CYP3A5 inhibitor / inducers taken while the patient is undergoing ADT may alter the efficacy of ADT. Although drug interactions monitor direct interactions between drugs the effect of concomitant CYP3A5 inducers and inhibitors have not been demonstrated previously. Based on our data we suggest that taking CYP3A5 inhibitors concomitantly may clinically benefit patients undergoing ADT (enhancing its effect), whereas taking CYP3A5 inducers may reduce the efficacy of the ADT treatment (countering its effect). These observations suggest that the effect of these inhibitors and inducers may be more relevant in AA patient as they tend to carry the wild type CYP3A5 and may result in therapeutic resistance. This study also suggests care be taken while prescribing CYP3A5 inducers when the patients are undergoing ADT. In addition, it also suggests that genetic testing for CYP3A5 polymorphism in patients may provide significant information about the potential impact of these interactions, facilitating personalized treatment regimens.

---

**Supplementary Table 1:**
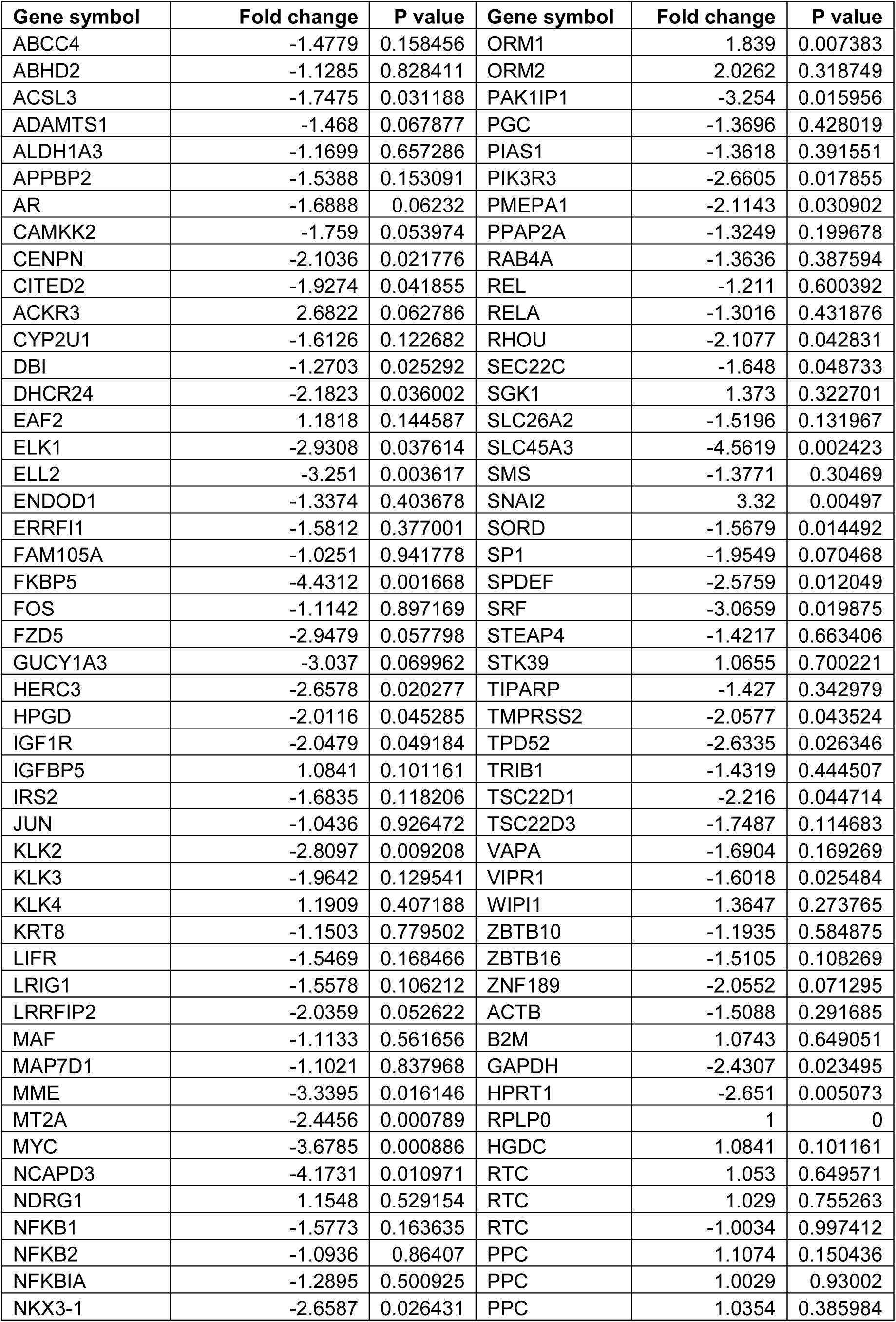
Table showing fold change with P value in AR downstream-regulated genes.

## References

1. Huggins C, Hodges CV: Studies on prostatic cancer. I. The effect of castration, of estrogen and of androgen injection on serum phosphatases in metastatic carcinoma of the prostate. 1941. J Urol 2002, 167(2 Pt 2):948–951; discussion 952.

2. Sadi MV, Walsh PC, Barrack ER: Immunohistochemical study of androgen receptors in metastatic prostate cancer. Comparison of receptor content and response to hormonal therapy. Cancer 1991, 67(12):3057–3064.

3. van der Kwast TH, Schalken J, Ruizeveld de Winter JA, van Vroonhoven CC, Mulder E, Boersma W, Trapman J: Androgen receptors in endocrine-therapy-resistant human prostate cancer. Int J Cancer 1991, 48(2):189–193.

4. Chodak GW, Kranc DM, Puy LA, Takeda H, Johnson K, Chang C: Nuclear localization of androgen receptor in heterogeneous samples of normal, hyperplastic and neoplastic human prostate. J Urol 1992, 147(3 Pt 2):798–803.

5. Heinlein CA, George Whipple Laboratory for Cancer Research DoP, Urology, and Radiation Oncology, and the Cancer Center, University of Rochester, Rochester, New York 14642, Chang C, George Whipple Laboratory for Cancer Research DoP, Urology, and Radiation Oncology, and the Cancer Center, University of Rochester, Rochester, New York 14642: Androgen Receptor in Prostate Cancer. Endocrine Reviews 2004, 25(2):276–308.

6. de Bono JS, Logothetis CJ, Molina A, Fizazi K, North S, Chu L, Chi KN, Jones RJ, Goodman OB, Jr., Saad F et al: Abiraterone and increased survival in metastatic prostate cancer. N Engl J Med 2011, 364(21):1995–2005.

7. Bianchini D, Lorente D, Rodriguez-Vida A, Omlin A, Pezaro C, Ferraldeschi R, Zivi A, Attard G, Chowdhury S, de Bono JS: Antitumour activity of enzalutamide (MDV3100) in patients with metastatic castration-resistant prostate cancer (CRPC) pre-treated with docetaxel and abiraterone. Eur J Cancer 2014, 50(1):78–84.

8. Darshan MS, Loftus MS, Thadani-Mulero M, Levy BP, Escuin D, Zhou XK, Gjyrezi A, Chanel-Vos C, Shen R, Tagawa ST et al: Taxane-induced blockade to nuclear accumulation of the androgen receptor predicts clinical responses in metastatic prostate cancer. Cancer Res 2011, 71(18):6019–6029.

9. Mezynski J, Pezaro C, Bianchini D, Zivi A, Sandhu S, Thompson E, Hunt J, Sheridan E, Baikady B, Sarvadikar A et al: Antitumour activity of docetaxel following treatment with the CYP17A1 inhibitor abiraterone: clinical evidence for cross-resistance? Ann Oncol 2012, 23(11):2943–2947.

10. Mitra R, Goodman OB: CYP3A5 regulates prostate cancer cell growth by facilitating nuclear translocation of AR. Prostate 2015, 75:527–538.

11. Zanger UM, Schwab M: Cytochrome P450 enzymes in drug metabolism: regulation of gene expression, enzyme activities, and impact of genetic variation. Pharmacol Ther 2013, 138(1):103–141.

12. Yamakoshi Y, Kishimoto T, Sugimura K, Kawashima H: Human prostate CYP3A5: identification of a unique 5’-untranslated sequence and characterization of purified recombinant protein. Biochem Biophys Res Commun 1999, 260(3):676–681.

13. Hou G, Zheng Y, Wei D, Li X, Wang F, Tian J, Zhang G, Yan F, Zhu Z, Meng P et al: Development and validation of a SEER-based prognostic nomogram for patients with bone metastatic prostate cancer. Medicine (Baltimore) 2019, 98(39):e17197.

14. Tompkins LM, Wallace AD: Mechanisms of cytochrome P450 induction. J Biochem Mol Toxicol 2007, 21(4):176–181.

15. Lolodi O, Wang YM, Wright WC, Chen T: Differential Regulation of CYP3A4 and CYP3A5 and its Implication in Drug Discovery. Curr Drug Metab 2017, 18(12):1095–1105.

16. Nem D, Baranyai D, Qiu H, Godtel-Armbrust U, Nestler S, Wojnowski L: Pregnane X receptor and yin yang 1 contribute to the differential tissue expression and induction of CYP3A5 and CYP3A4. PLoS One 2012, 7(1):e30895.

17. Finnstrom N, Bjelfman C, Soderstrom TG, Smith G, Egevad L, Norlen BJ, Wolf CR, Rane A: Detection of cytochrome P450 mRNA transcripts in prostate samples by RT-PCR. Eur J Clin Invest 2001, 31(10):880–886.

18. Leskela S, Honrado E, Montero-Conde C, Landa I, Cascon A, Leton R, Talavera P, Cozar JM, Concha A, Robledo M et al: Cytochrome P450 3A5 is highly expressed in normal prostate cells but absent in prostate cancer. Endocr Relat Cancer 2007, 14(3):645–654.

19. Moilanen AM, Hakkola J, Vaarala MH, Kauppila S, Hirvikoski P, Vuoristo JT, Edwards RJ, Paavonen TK: Characterization of androgen-regulated expression of CYP3A5 in human prostate. Carcinogenesis 2007, 28(5):916–921.

20. Kuehl P, Zhang J, Lin Y, Lamba J, Assem M, Schuetz J, Watkins PB, Daly A, Wrighton SA, Hall SD et al: Sequence diversity in CYP3A promoters and characterization of the genetic basis of polymorphic CYP3A5 expression. Nat Genet 2001, 27(4):383–391.

21. Theodore S, Sharp S, Zhou J, Turner T, Li H, Miki J, Ji Y, Patel V, Yates C, Rhim JS: Establishment and characterization of a pair of non-malignant and malignant tumor derived cell lines from an African American prostate cancer patient. Int J Oncol 2010, 37(6):1477–1482.

22. Hooker SE, Woods-Burnham L, Bathina M, Lloyd SM, Gorjala P, Mitra R, Nonn L, Kimbro KS, Kittles R: Genetic ancestry analysis reveals misclassification of commonly used cancer cell lines. Cancer Epidemiol Biomarkers Prev 2019.

23. Lynch T, Price A: The effect of cytochrome P450 metabolism on drug response, interactions, and adverse effects. Am Fam Physician 2007, 76(3):391–396.

24. Schuetz EG, Relling MV, Kishi S, Yang W, Das S, Chen P, Cook EH, Rosner GL, Pui CH, Blanco JG et al: PharmGKB update: II. CYP3A5, cytochrome P450, family 3, subfamily A, polypeptide 5. Pharmacol Rev 2004, 56(2):159.

25. Lamba JK, Lin YS, Schuetz EG, Thummel KE: Genetic contribution to variable human CYP3A-mediated metabolism. Adv Drug Deliv Rev 2002, 54(10):1271–1294.

26. Lamba J, Hebert JM, Schuetz EG, Klein TE, Altman RB: PharmGKB summary: very important pharmacogene information for CYP3A5. Pharmacogenet Genomics 2012, 22(7):555–558.

27. Xu J, Kalos M, Stolk JA, Zasloff EJ, Zhang X, Houghton RL, Filho AM, Nolasco M, Badaro R, Reed SG: Identification and characterization of prostein, a novel prostate-specific protein. Cancer Res 2001, 61(4):1563–1568.

28. Kalos M, Askaa J, Hylander BL, Repasky EA, Cai F, Vedvick T, Reed SG, Wright GL, Jr., Fanger GR: Prostein expression is highly restricted to normal and malignant prostate tissues. Prostate 2004, 60(3):246–256.

29. Avellino R, Romano S, Parasole R, Bisogni R, Lamberti A, Poggi V, Venuta S, Romano MF: Rapamycin stimulates apoptosis of childhood acute lymphoblastic leukemia cells. Blood 2005, 106(4):1400–1406.

30. Pei H, Li L, Fridley BL, Jenkins GD, Kalari KR, Lingle W, Petersen G, Lou Z, Wang L: FKBP51 affects cancer cell response to chemotherapy by negatively regulating Akt. Cancer Cell 2009, 16(3):259–266.

31. Bolton EC, So AY, Chaivorapol C, Haqq CM, Li H, Yamamoto KR: Cell- and gene-specific regulation of primary target genes by the androgen receptor. Genes Dev 2007, 21(16):2005–2017.

32. Ni L, Yang CS, Gioeli D, Frierson H, Toft DO, Paschal BM: FKBP51 promotes assembly of the Hsp90 chaperone complex and regulates androgen receptor signaling in prostate cancer cells. Mol Cell Biol 2010, 30(5):1243–1253.

33. Grad JM, Dai JL, Wu S, Burnstein KL: Multiple androgen response elements and a Myc consensus site in the androgen receptor (AR) coding region are involved in androgen-mediated up-regulation of AR messenger RNA. Mol Endocrinol 1999, 13(11):1896–1911.

34. Koh CM, Bieberich CJ, Dang CV, Nelson WG, Yegnasubramanian S, De Marzo AM: MYC and Prostate Cancer. Genes Cancer 2010, 1(6):617–628.

35. Tomlins SA, Mehra R, Rhodes DR, Cao X, Wang L, Dhanasekaran SM, Kalyana-Sundaram S, Wei JT, Rubin MA, Pienta KJ et al: Integrative molecular concept modeling of prostate cancer progression. Nat Genet 2007, 39(1):41–51.

36. Bai S, Cao S, Jin L, Kobelski M, Schouest B, Wang X, Ungerleider N, Baddoo M, Zhang W, Corey E et al: A positive role of c-Myc in regulating androgen receptor and its splice variants in prostate cancer. Oncogene 2019.

37. Gao L, Schwartzman J, Gibbs A, Lisac R, Kleinschmidt R, Wilmot B, Bottomly D, Coleman I, Nelson P, McWeeney S et al: Androgen receptor promotes ligand-independent prostate cancer progression through c-Myc upregulation. PLoS One 2013, 8(5):e63563.

38. Qiu X, Pascal LE, Song Q, Zang Y, Ai J, O’Malley KJ, Nelson JB, Wang Z: Physical and Functional Interactions between ELL2 and RB in the Suppression of Prostate Cancer Cell Proliferation, Migration, and Invasion. Neoplasia 2017, 19(3):207–215.

39. Sims RJ, 3rd, Belotserkovskaya R, Reinberg D: Elongation by RNA polymerase II: the short and long of it. Genes Dev 2004, 18(20):2437–2468.

40. Hieronymus H, Lamb J, Ross KN, Peng XP, Clement C, Rodina A, Nieto M, Du J, Stegmaier K, Raj SM et al: Gene expression signature-based chemical genomic prediction identifies a novel class of HSP90 pathway modulators. Cancer Cell 2006, 10(4):321–330.

41. Pascal LE, Masoodi KZ, Liu J, Qiu X, Song Q, Wang Y, Zang Y, Yang T, Wang Y, Rigatti LH et al: Conditional deletion of ELL2 induces murine prostate intraepithelial neoplasia. J Endocrinol 2017, 235(2):123–136.

42. Zhang W, Liu B, Wu W, Li L, Broom BM, Basourakos SP, Korentzelos D, Luan Y, Wang J, Yang G et al: Targeting the MYCN-PARP-DNA Damage Response Pathway in Neuroendocrine Prostate Cancer. Clin Cancer Res 2018, 24(3):696–707.

43. Kumar A, Mikolajczyk SD, Goel AS, Millar LS, Saedi MS: Expression of pro form of prostate-specific antigen by mammalian cells and its conversion to mature, active form by human kallikrein 2. Cancer Res 1997, 57(15):3111–3114.

44. Haese A, Graefen M, Becker C, Noldus J, Katz J, Cagiannos I, Kattan M, Scardino PT, Huland E, Huland H et al: The role of human glandular kallikrein 2 for prediction of pathologically organ confined prostate cancer. Prostate 2003, 54(3):181–186.

45. Stephan C, Jung K, Nakamura T, Yousef GM, Kristiansen G, Diamandis EP: Serum human glandular kallikrein 2 (hK2) for distinguishing stage and grade of prostate cancer. Int J Urol 2006, 13(3):238–243.

46. Steuber T, Vickers AJ, Serio AM, Vaisanen V, Haese A, Pettersson K, Eastham JA, Scardino PT, Huland H, Lilja H: Comparison of free and total forms of serum human kallikrein 2 and prostate-specific antigen for prediction of locally advanced and recurrent prostate cancer. Clin Chem 2007, 53(2):233–240.

47. Shang Z, Niu Y, Cai Q, Chen J, Tian J, Yeh S, Lai KP, Chang C: Human kallikrein 2 (KLK2) promotes prostate cancer cell growth via function as a modulator to promote the ARA70-enhanced androgen receptor transactivation. Tumour Biol 2014, 35(3):1881–1890.

